# Targeting cellular metabolism to inhibit synergistic biofilm formation of multi-species isolated from a cooling water system

**DOI:** 10.1101/2021.01.28.428600

**Authors:** Dingrong Kang, Wenzheng Liu, Fatemeh Bajoul kakahi, Frank Delvigne

## Abstract

Biofilm is ubiquitous in natural environments, causing biofouling in industrial water systems and leading to liquidity and heat transfer efficiency decreases. In particular, multi-species coexistence in biofilms can provide the synergy needed to boost biomass production and enhance treatment resistance. In this study, a total of 37 bacterial strains were isolated from a cooling tower where acetic acid and propionic acid were used as the primary carbon sources. These isolates mainly belonged to Proteobacteria and Firmicutes, which occupied more than 80% of the total strains according to the 16S rRNA gene amplicon sequencing. Four species (*Acinetobacter* sp. CTS3*, Corynebacterium* sp. CTS5*, Providencia* sp. CTS12, and *Pseudomonas* sp. CTS17) were observed to co-exist in the synthetic medium, showing a synergistic effect towards biofilm formation. Three metabolic inhibitors (sulfathiazole, 3-Bromopyruvic acid, and 3-Nitropropionic acid) were employed as possible treatments against biofilm formation due to their inhibition effect on c-di-GMP biosynthesis or assimilation of volatile fatty acids. All of them displayed evident inhibition profiles to biofilm formation. Notably, the combination of these three inhibitors possessed a remarkable ability to block the development of a multi-species biofilm with lower concentrations, suggesting an enhanced effect with their simultaneous use. This study demonstrates that targeting cellular metabolism is an effective way to inhibit biofilm formation derived from multi-species.

## 1. Introduction

Diverse microorganisms frequently co-exist in natural environments as biofilms, producing extracellular polymeric substances that play essential roles in establishing spatial structure and generating new properties (Elias and Banin, 2012; Karygianni et al., 2020). The emerging biofilm properties contribute to improving nutrient acquisition, enhancing antibiotic tolerance, and strengthening internal cooperation towards resisting attacks or stresses from the external environment (Flemming et al., 2016). However, microbial biofilm formation phenomena can cause serious biofouling with concomitant efficiency reductions or failure in industrial water-using systems, such as cooling towers (Bott, 1998). The latter industrial system’s water sources are commonly derived from groundwater and surface waters (AM Peer and T Sanders, 2016). Recently, there has been an increasing interest in using municipal or plant wastewater to replace freshwater consumption as it is environmentally beneficial and is easier to ensure abundant supply and sustainable development (Li et al., 2011). However, wastewater has abundant nutrient substances, such as volatile fatty acids (acetate, propionate, and butyrate), which are essential intermediates in microbial fermentation processes, resulting in plentiful microorganisms capable of growing and forming biofilm (Zhou et al., 2018). Therefore, seeking an effective strategy to reduce biofilm formation resulting from wastewater has been an enticing research area to prevent biofouling in industrial water systems.

Microbial communities in cooling towers are highly dynamic, changing within seasons and locations due to the unique environmental characteristics; albeit with their differences, they generally share taxa associated with biofilm formation (Tsao et al., 2019). A common core microbiome was identified from biofilms of four cooling towers filled with different water sources (Di Gregorio et al., 2017). In particular, *Pseudomonas* was acknowledged to be within the pioneer colonizers and then followed by multiple species able to co-colonize and form mixed biofilms (Doğruöz et al., 2009). Following the interspecific interactions, including cross-feeding and metabolic exchange among mixed biofilms, facilitate the microbes to survive in harsh environments, enhance biomass production, and increase resistance to external stress (Tan et al., 2017). Therefore, targeting key species related to biofilm development rather than the free-living cells has been found effective in blocking the mixed-species biofilm formation.

Several approaches have been developed to limit mixed-species biofilm formation in cooling water systems. The most frequent countermeasure used is by addition of oxidizing or non-oxidizing biocides, targeting essential components such as the cellular walls, membranes, structural proteins, and RNA/DNA of microorganisms, killing the microbial cells (Di Pippo et al., 2018). However, microbes residing in biofilms display higher tolerance to these biocides in comparison to planktonic cells, and their resistance in biofilms can increase during prolonged treatment regimes through phenotypic diversification and/or horizontal gene transfers (Fux et al., 2005; Harrison et al., 2007). Material modifications such as coatings with silver and copper nanoparticles have shown their efficacy to prevent the initial biofilm attachment in the cooling pipes (Ogawa et al., 2016). Imidazolium and piperidinium-based ionic liquids were also found to inhibit cell adhesion to the surfaces of different materials, leading to blocked biofilm formation (Anandkumar et al., 2020; Reddy et al., 2017). Furthermore, using the quorum quenching bacteria (Jo et al., 2016), ultrasonic treatments (Rodríguez-Calvo et al., 2020), and electromagnetic processes (Xiao et al., 2020) have been explored towards obstructing biofilm development or eliminating the formed biofilms. Despite all these diverse biofilms contending strategies, few of them have been implemented with great success to field antifouling applications due to many limitations such as strain specificity, stability, and long-term efficacy (Flemming, 2020).

Metabolic inhibitors have been applied to target specific microorganism pathways, resulting in diverse gene expression regulative cascades and the concomitant reduction or loss of distinct metabolic cell functions (Mohiuddin et al., 2020; Saiardi et al., 2018). Therefore, different metabolic inhibitors could be used to inhibit specific phenotypical traits, such as biofilm formation (Mulhbacher et al., 2010; Wan et al., 2019). Regarding the latter, various metabolic pathways have been found to associate with biofilm development in recent years (Armbruster and Parsek, 2018; Rabin et al., 2015). For example, previous studies have illustrated that second messenger cyclic di-GMP (c-di-GMP) guides the bacterial switching behavior from a planktonic state to a biofilm formation lifestyle (Valentini and Filloux, 2016). Inhibiting the synthesis pathway of c-di-GMP was found to be an effective means to control biofilm formation (Sambanthamoorthy et al., 2012). Specifically, sulfathiazole (ST) has been reported to interfere with c-di-GMP metabolism, leading to efficiently inhibiting biofilm formation (Antoniani et al., 2010). Additionally, 3-Bromopyruvic acid (3BP) and 3-Nitropropionic acid (3-NP) have been found to block the assimilation of volatile fatty acids in cellular metabolism through the inactivation of key enzymes (Sharma et al., 2000). In this study, we studied the possibility of biofilm formation inhibition on industrial water systems by using three different metabolic inhibitors while testing their efficacy and specificity. Firstly, cultivable bacteria were isolated and identified from a cooling tower. Representative strains were then used to screen and characterize synergistic biofilm community. Subsequently, ST, 3BP, and 3-NP were employed to inhibit metabolic pathways related to c-di-GMP biosynthesis and/or assimilation of volatile fatty acids, and its biofilm formation inhibition was measured. The present work pretends to contribute towards constructing an alternative strategy to overcome biofouling by targeting the cellular metabolism associated with biofilm formation.

## 2. Material and methods

### 2.1 Biofilm sampling and synthetic medium

Biofilm sample was collected from a cooling tower of a French industry plant that produces sugar and alcohol. The cooling tower was fed by wastewater with a pH range from 7.0 to 9.0, at a temperature of 28 °C. The main carbon sources found in wastewater were volatile fatty acids, including propionic acid (1.50 g/L), acetic acid (1.00 g/L), lactic acid (0.016 g/L), and formic acid (0.013 g/L). To reproduce the industrial conditions, a mimetic cooling tower environment was lab operated using a synthetic medium. Synthetic medium was based on M9 minimal medium (33.7 mM Na_2_HPO_4_, 22.0 mM KH_2_PO_4_, 8.55 mM NaCl, 9.35 mM NH_4_Cl, 1 mM MgSO_4_, 0.3 mM CaCl_2_, trace elements solution (13.4 mM EDTA, 3.1 mM FeCl_3_-6H_2_O, 0.62 mM ZnCl_2_, 76 μM CuCl_2_-2H_2_O, 42 μM CoCl_2_-2H_2_O, 162 μM H_3_BO_3_, 8.1 μM MnCl_2_-4H_2_O), 1 μg/L biotin and 1 μg/L thiamin), and supplemented with propionic acid (1.50 g/L) and acetic acid (1.00 g/L) as main carbon sources (PH = 7.2).

### 2.2 Strains isolation from biofilm sample

A total of 10 mL of biofilm sample (three biological replicates) were placed into a 50 ml centrifuge tube containing 15 sterile glass beads (diameter 2 mm) and an equal volume of sterile phosphate-buffered saline solution (PBS). The sample was then homogenized using a vortex mixer for 30 s at 2500 rpm and serially diluted to 1 × 10^-7^ with PBS. A 100 μL aliquot of each serial dilution was spread onto LB agar plates in triplicate. Agar plates were incubated at 30 °C for two to five days. Colonies were selected by differentiable morphologies.

### 2.3 Identification of isolated strains

Obtained strains were grown in fresh LB medium overnight and spread on LB plates three times to ensure pure cultures. 2 mL suspension of each pure culture was then used for DNA extraction. DNA was extracted by using Genejet Genomic DNA Purification Kit according to the manufacturer’s instructions. Following DNA was used as a template to amplify and sequence the full 16S rRNA gene with primers (8F 5’ - AGAGTTTGATCCTGGCTCAG - 3’; 1492R 5’ - ACGGTTACCTTGTTACGACTT - 3’) by Sanger method. Full sequences were checked and jointed manually using 4Peaks software (https://nucleobytes.com/4peaks/index.html). The Resulting 16S rRNA gene sequences were queried and identified with reference sequences in the Silva database. Sequencing data are available at the NCBI GenBank database with the accession numbers MW389057 – MW389093, including *Acinetobacter* sp. CTS3 (MW389059), *Corynebacterium* sp. CTS5 (MW389060), *Providencia* sp. CTS12 (MW389066), and *Pseudomonas* sp. CTS17 (MW389070).

### 2.4 Constructing the phylogenetic trees of isolated strains

For constructing the phylogenetic tree for the total isolates obtained from the cooling tower, obtained 16s rRNA gene sequences were aligned and processed in MEGAX software (Kumar et al., 2018). The evolutionary relationship was inferred by using the Maximum Likelihood method and the Tamura-Nei model. To uncover the phylogenetic evolution relationship of single strains within the genera, 16S rRNA genes of reference strains were downloaded from the NCBI database and aligned with the resulting sequences. The phylogenetic trees of each single strain were constructed by using the process described above.

### 2.5 Screening multi-species biofilms formation by using crystal violet (CV) assay

Ten representative strains from four phyla were selected for the co-existence assay, which was performed with ten successive cycles of repeated batch cultivation using synthetic medium. The purpose of this assay was to further screen multi-species with potential cooperative capabilities for biofilm formation in the cooling tower. In the end, four strains were obtained. Then they were grown as single- and multi-species biofilms using a Nunc-TSP lid culture system which comprises a 96-well plate lid with pegs extending into each well. Biofilm formation was quantified by crystal violet (CV) staining after 24 h and 48 h of cultivation. Pre-cultures were grown overnight up to OD_600nm_ 1.0 at 30 °C. The cell suspensions were then adjusted to 10^8^ cells per milliliter in the synthetic medium. A total of 160 μL monocultures or mixed cultures (equal volume for each) were added to each well. Fresh medium was used as the negative control. The plates were sealed with Parafilm and incubated with shaking 200 rpm at 30 °C. The biofilm biomass was measured as previously reported CV assays (Ren et al., 2014). CV quantification was performed on the pegs of the Nunc-TSP lid culture system. The peg lids were taken out and washed three times using PBS after 24 h and 48 h of cultivation. Subsequently, the peg lids were placed in plates with 180 μL of an aqueous 1% (w/v) CV solution. The lids were rewashed with PBS three times after staining for 20 min. Peg lids were put into a new microtiter plate with 200 μL of 33% glacial acetic acid in each well. After 15 min, the dissolved CV solution’s absorbance was determined at 590 nm using a microplate reader (Tecan, Spark).

### 2.6 Inhibition assay of cell proliferation

The cell suspensions of four strains were adjusted to 10^8^ cells per milliliter in the synthetic medium separately. A total of 200 μL monocultures or mixed cultures (equal volume for each) were added to each well of 96-well plates. Different final concentrations of metabolic inhibitors (ST, 3BP, and 3-NP) from 2 μg/mL to 256 μg/mL were supplied to the bacterial suspension. No inhibitor addition treatment was used as control. Then the plates with cultures were placed on the shaker with 200 rpm at 30 °C. Cell proliferation was measured based on the net increase of the OD_600nm_ using a microplate reader (Tecan, Spark) after 24 h and 48 h of cultivation.

### 2.7 Inhibition assay of biofilm formation

A total of 160 μL of bacterial suspension (1×10^8^ cells/mL) in terms of monoculture and mixed cultures were prepared as above, then added to the Nunc-TSP lid culture system. Following metabolic inhibitors and their combinations were supplied to the suspension with different final concentrations from 2 μg/mL to 256 μg/mL, with no inhibitor addition as the control. The plates were then placed on the shaker at 200 rpm and 30 °C. The biofilm quantification assays were performed as described above using the CV staining assay after 24 h and 48 h cultivation. The CV absorbance at 590 nm was used to evaluate the inhibiting effect of the metabolic inhibitors on biofilm formation.

### 2.8 Statistical Analysis

All of the experiments were performed at least in four biological replicates. One-way ANOVA followed by post hoc Tukey’s HSD tests were used for establishing the statistical differences of biofilm-forming capabilities. The significance level was set to *p* < 0.05.

## 3. Results

### 3.1 Isolation and identification of strains from biofilm

The viable strains that displayed distinct morphological features were isolated and identified from the cooling tower biofilm samples to characterize the cultivable microbial composition. Thirty-seven strains were found to belong to four different phyla: Actinobacteria, Proteobacteria, Bacteroidetes, and Firmicutes (Fig. 1). Twenty strains were identified as Proteobacteria consisting of nine genera, of which *Pseudomonas* was the most extensive branch. To further obtain a simplified microbial community with the capacity to use volatile fatty acids collectively, ten representative strains from the four phyla were selected to co-culture in a continuous mode. A total of four species, including *Acinetobacter* sp. CTS3 (A3)*, Corynebacterium* sp. CTS5 (C5)*, Providencia* sp. CTS12 (P12), and *Pseudomonas* sp. CTS17 (P17) were verified to be persistent. Phylogenic trees were built to explore their phylogenic relationship between the different species (Fig. S1-4). The result shows that the closest evolutionary relationship for the A3 strain is *Acinetobacter johnsonii* (Fig. S1). Similarly, *Pseudomonas composti* was found to be the closest related species for the P17 strain (Fig. S4). For C5 and P12, the closest references are *Corynebacterium glutamicum* and *Providencia heimbachae*, respectively (Fig. S2 and S3).

**Fig. 1.**
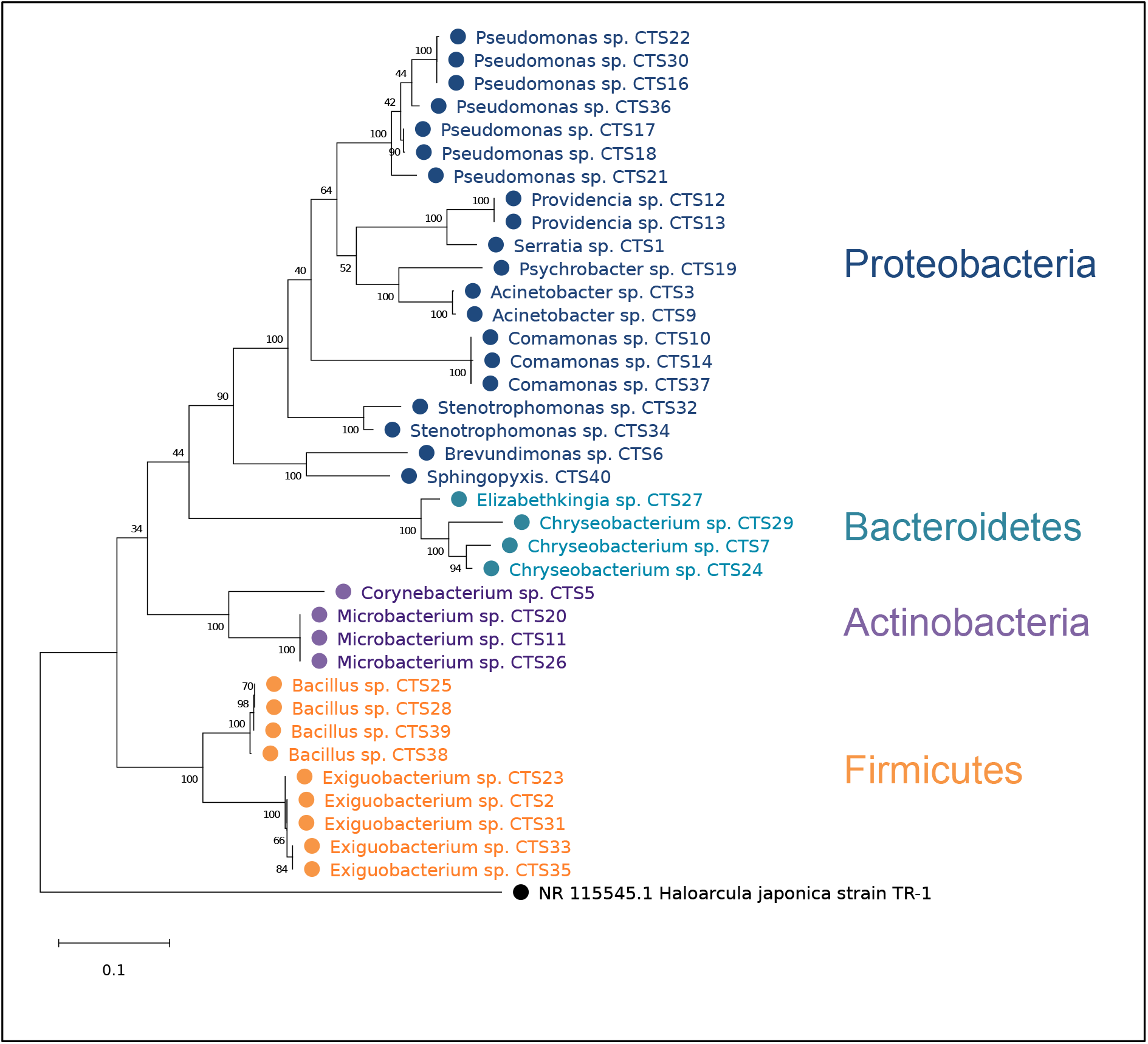
Phylogenetic tree of the strains isolated from biofilm sample. *Halocarcula japonica* strain TR-1 (GenBank: NR115545.1) was set as the outgroup species. Bootstrap values are displayed at each node.

### 3.2 Designing multi-species biofilms with synergistic interactions

To obtain the multi-species biofilms and characterize their possible synergetic effect, all potential combinations of the four strains (single, dual-, triple-, and quadruple species) were cultured in the synthetic medium (Fig. 2). Suspended cell density and biofilm-forming capacity were measured after 24 h and 48 h. Four strains were found with different cell densities (OD600nm) after the cultivation, suggesting that they can grow independently but exhibit different capacities to assimilate the volatile fatty acids. For all mixture cultivations, the A3P17 pair performed with the highest cell density among all other combinations with an OD600nm value of 0.65 after 24 h. The other mixtures’ cell density (OD_600nm_) range was found to be between 0.24 and 0.63 after 24 h. A3, C5, and P12 single strain cultures showed a low capacity to form biofilm, with CV values (Abs_590nm_) below 0.25. In contrast, the CV value (Abs_590nm_) from P17 culture showed a higher than 10-fold increase displaying a strong capacity for biofilm production. It is worth noting that all of the bacterial combinations comprising P17 possessed a strong ability to form biofilm. The latter was predominantly observed in the four-species combination, in which the CV value (Abs_590nm_) reached up to 8.30 after 24 h, suggesting an evident synergistic effect on biofilm formation among these species.

**Fig. 2.**
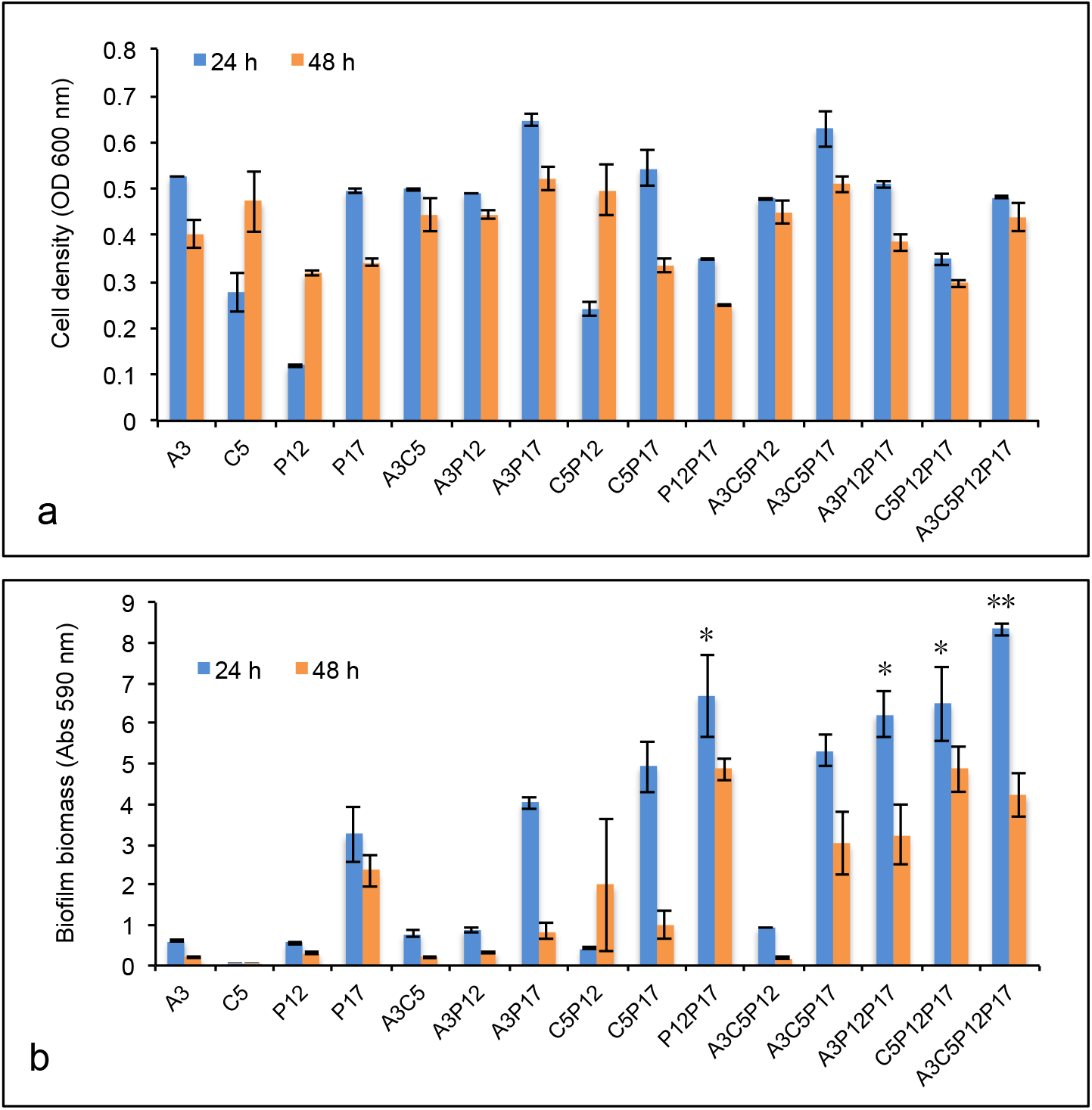
Comparison of cell density and biofilm biomass from single and multiple species in the synthetic medium after 24 h and 48 h. (a). Cell density. (b). Biofilm biomass, n ≥ 6. The significance level of biofilm formation after 24 h between P17 and multi-species comprising P17 was shown by “*” (*p* < 0.05) and “**” (*p* < 0.001).

### 3.3 Inhibitory effect of ST, 3BP, and 3-NP on cell proliferation

Three chemical agents, i.e., sulfathiazole (ST), 3-Bromopyruvic acid (3BP), and 3-Nitropropionic acid (3-NP), were applied as the metabolic inhibitors to act on the cell proliferation of mono- and multi-species (four species). Cell density was recorded within the treatment of different concentrations after 24 h and 48 h cultivation (Fig. 3). Three agents displayed different inhibitory effects on cell proliferation among four mono-species. ST cannot inhibit any of them entirely below the concentration of 256 μg/mL, but it showed a visible inhibiting effect to P17, especially as the concentration above 64 μg/mL. 3BP was able to prevent the cell proliferation of P12 and P17 almost entirely after 24 h with a concentration of 256 μg/mL, following the inhibiting effect was still acting on P17, but not P12 after 48 h. Notably, 3-NP presented a more substantial inhibitory effect than the other two (ST, 3BP) on these four species. The cell proliferation of C5 and P12 was inhibited completely with concentrations of 2 μg/mL and 8 μg/mL, respectively. Additionally, cell proliferation of multi-species showed a distinct performance with these three metabolic inhibitors. Cell density was reduced along with increasing concentrations of ST and 3BP after 24 h, while it was not affected even higher than the control after 48 h. Nevertheless, 3-NP still displayed the effective inhibition to the multi-species in 24 h (64 μg/mL) and 48 h (256 μg/mL), which was able to inhibit them completely.

**Fig. 3.**
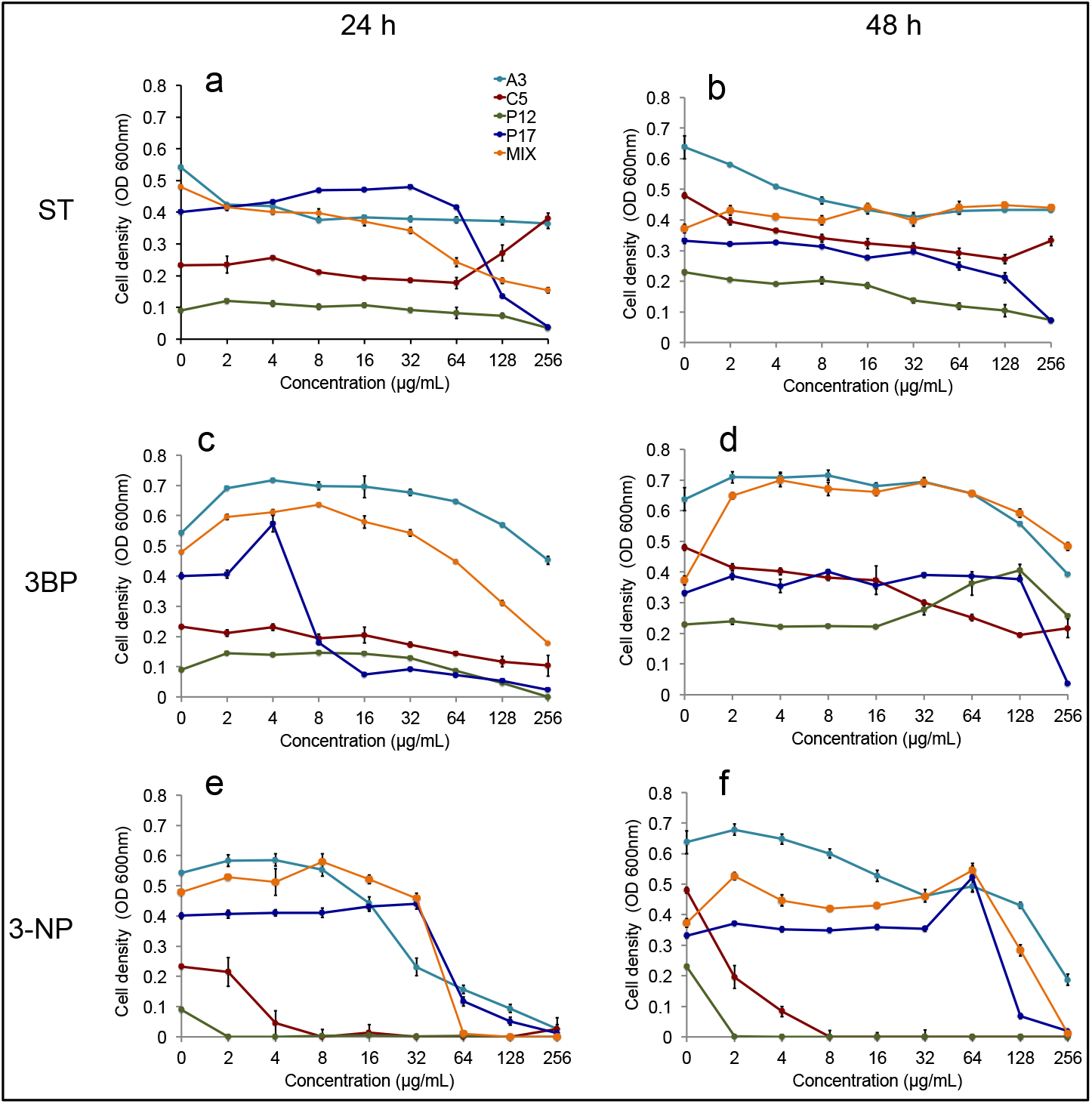
Cell density of P17 and four species (MIX) with different metabolic inhibitors in the synthetic medium after 24 h and 48 h. ST: sulfathiazole, 3BP: 3-Bromopyruvic acid, and 3-NP: 3-Nitropropionic acid, n = 4.

### 3.4 Inhibitory effect of ST, 3BP, and 3-NP on biofilm formation

Serial concentrations of ST, 3BP, and 3-NP were added to the Nunc-TSP lid system for P17 and multi-species (four species) cultures to evaluate the inhibitory effect on biofilm formation (Fig. 4). These three inhibitors displayed different inhibition patterns on the biofilm-forming capacity of P17. This inhibitory effect was proved to increase along with increasing inhibitor concentrations, from 64 μg/mL up to 256 μg/mL. On the other hand, low inhibitor concentrations seem to boost biofilm formation, especially with 3BP. Notably, more than ninety-five percentual reduction on biofilm formation were achieved with 3BP after 24 h at a concentration of 16 μg/mL and at 48 h with a concentration of 32 μg/mL. 3-NP and ST also were able to inhibit the biofilm formation effectively within concentrations of 64 μg/mL and 128 μg/mL. The three inhibitors also displayed an inhibitory effect on the multi-species culturing. 3-NP showed the highest inhibitory capacity even at low concentrations of 2 μg/mL after 24 h, while biofilm was formed after 48 h. Similarly, multi-species biofilms were enhanced after 48 h compared to the biofilm biomass quantification at 24 h with the other two inhibitors. Despite the latter, when the inhibitor concentration reached 256 μg/mL, biofilm formation was completely inhibited after 24 h and presented a more than 95% reduction after 48 h.

**Fig. 4.**
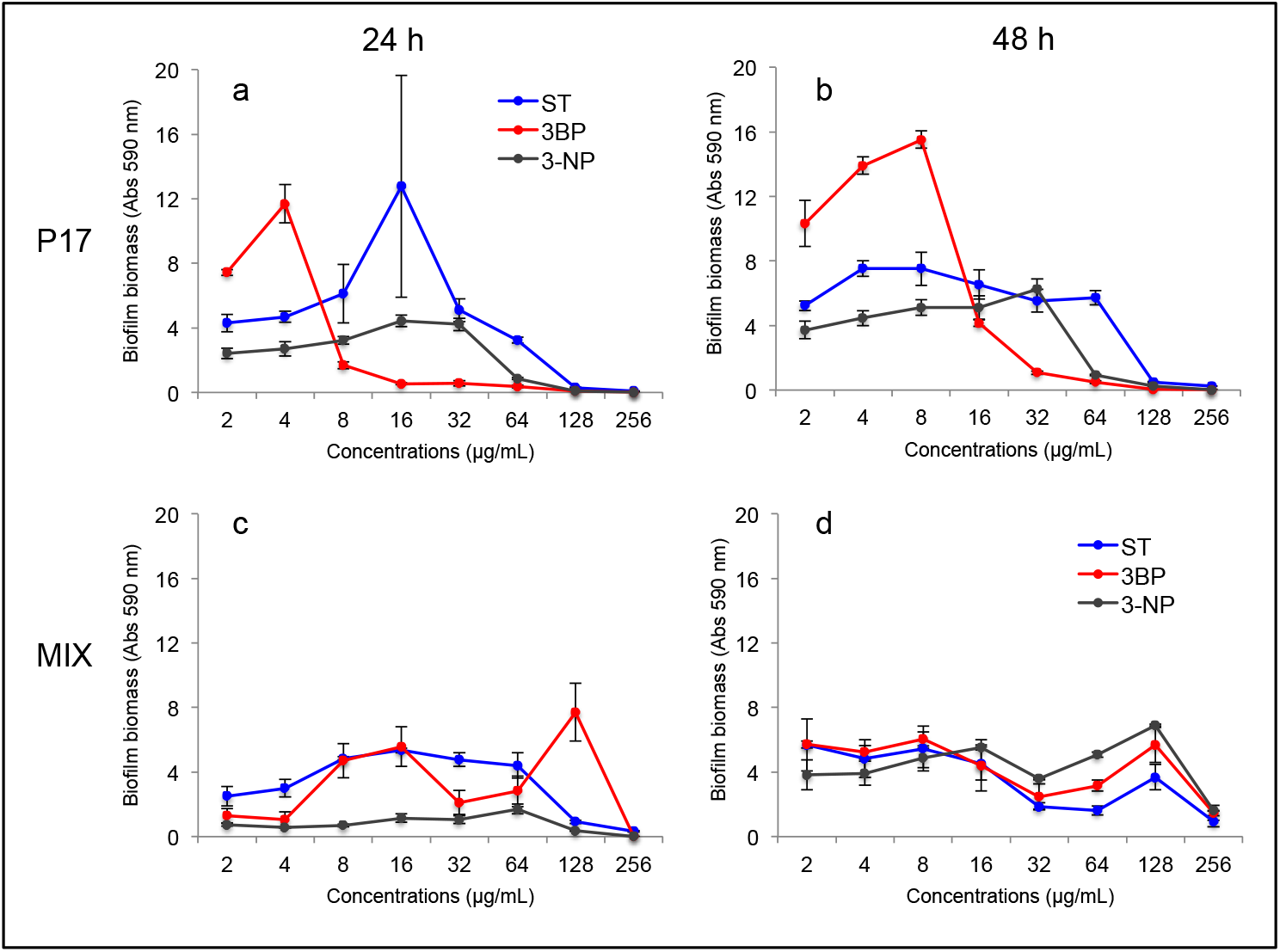
Biofilm formation of P17 and four species (MIX) with different metabolic inhibitors in the synthetic medium after 24 h and 48 h. ST: sulfathiazole, 3BP: 3-Bromopyruvic acid, and 3-NP: 3-Nitropropionic acid, n = 4.

### 3.5 Enhancement of inhibitory effect on biofilm formation *via* inhibitor combinations

Additionally, combinations of ST, 3BP, and 3-NP were used against the mono- and multi-species to investigate these molecules’ possible enhancement effect in preventing biofilm formation. All of the inhibitor combinations showed a remarkable inhibitory effect on biofilm formation in P17 and multi-species (four species) cultures (Fig. 5). Dual inhibitors combination (ST and 3BP) inhibited P17 biofilm formation almost wholly at a concentration of 8 μg/mL within 48 h. These results show a remarkably higher inhibitory effect from simultaneous use of ST and 3BP compared to their independent usage (ST: 128 μg/mL, 3BP: 64 μg/mL). Similarly, the other dual-inhibitor combinations (ST:3-NP and 3BP:3-NP) also presented strong inhibitory effects on biofilm formation. They were able to inhibit the biofilm-forming of P17 and multi-species entirely with a concentration of 64 μg/mL after 24 h and 48 h, of which the concentration value was found to be lower than their individual usage. Notably, the combination of the three inhibitors displayed an extreme inhibitory efficiency on the biofilm formation, which inhibited the biofilm-forming of P17 and multi-species entirely by treating with concentrations as low as 4 μg/mL and 8 μg/mL, respectively. Only low biofilm formation was detected even after 96 h; it showed that the combination was able to impact the biofilm-forming persistently (Fig. S5). These results demonstrate that an enhancement effect exists among the inhibitors, especially for combining three of them. The latter and their inhibition characterization make it clear that they could be part of a promising regulated strategy to biofilm formation of mono- and multi-species.

**Fig. 5.**
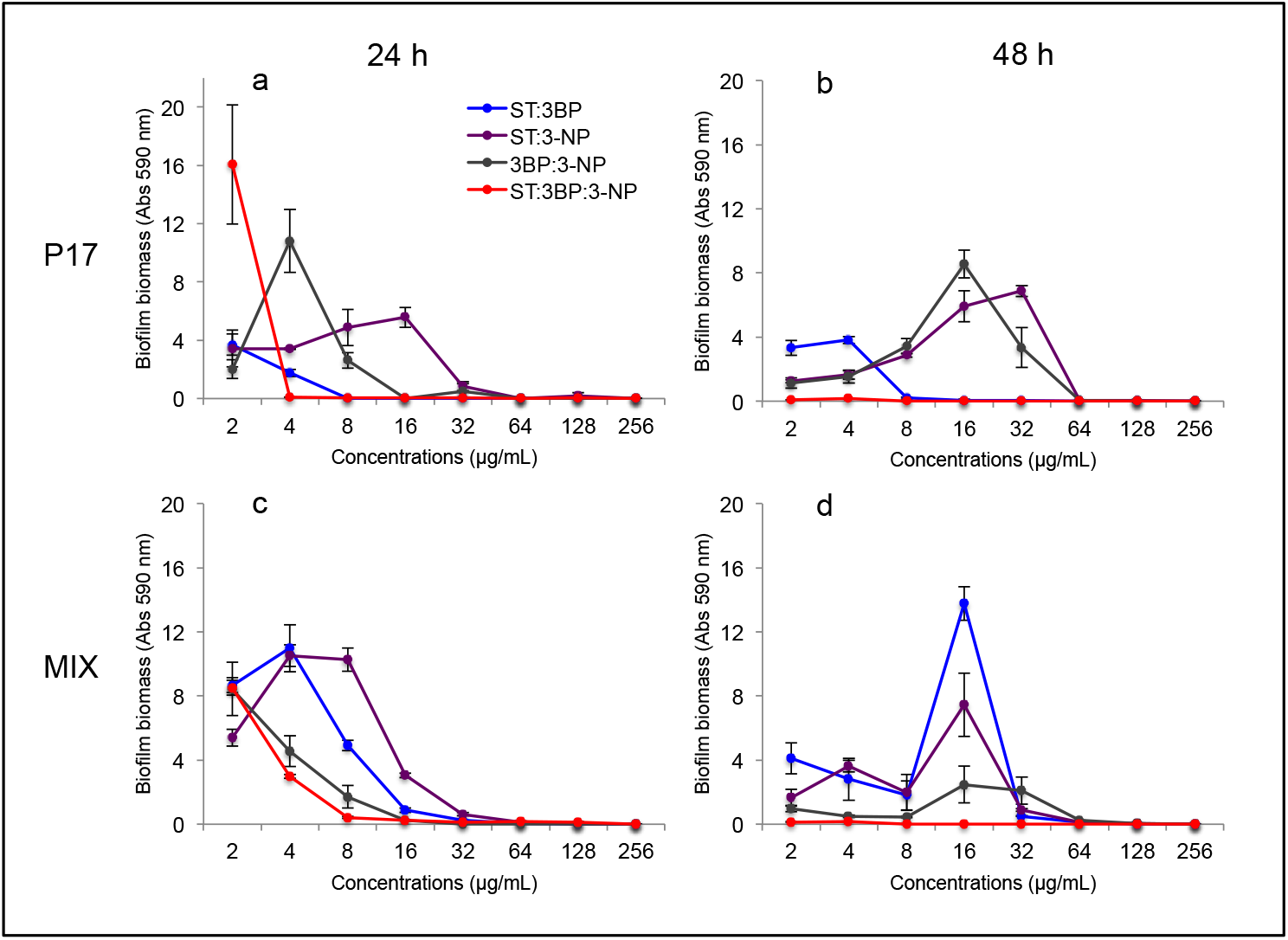
Biofilm formation of P17 and four species (MIX) with treatment of metabolic inhibitor combinations in the synthetic medium after 24 h and 48 h. ST: sulfathiazole, 3BP: 3-Bromopyruvic acid, and 3-NP: 3-Nitropropionic acid, n = 4.

## 4. Discussion

In this study, cultivable bacterial isolates were identified from biofilm samples originated from a cooling tower. The capacity of biofilm formation among representative species was evaluated, showing the synergistic biofilm formation effect among the four species in synthetic medium. Three metabolic inhibitors (ST, 3BP, and 3-NP) were selected against cell proliferation and biofilm formation owed to their ability to target the vital metabolic pathways in cellular metabolism. They displayed distinct degrees of inhibition effect to the cell proliferation of these four species, and all of them can inhibit the biofilm development of P17 and multi-species individually with concentrations lower than 256 μg/mL. Notably, the combination of three inhbitors possessed potent inhibition against the mono- and multi-species biofilms in a long term, employed with a much low concentration (8 μg/mL) from each. Targeting cellular metabolism via distinct functional metabolic inhibitors turns into a promising approach to prevent synergistic biofilm formation of multi-species.

Many studies have reported the microbial composition in the suspension of the cooling water systems. A combined NGS-based approach was used to study the whole bacterial community from a cooling tower, which displayed a highly diverse bacterial community conformed of more than 808 genera observed (Pereira et al., 2017). Seven species producing extracellular polymeric substances were isolated and characterized from the cooling tower through sequential culturing in glucose, sucrose, or galactose enriched medium (Ceyhan and Ozdemir, 2008). Here, 37 bacterial strains were identified in the cooling water biofilm mainly belonged to Proteobacteria and Firmicutes, which occupied more than 80% of the total isolates. Proteobacteria has already been reported to be the predominant bacteria in biofilms growing on recirculating cooling-water systems (MacDonald and Brözel, 2000; Pinel et al., 2020). Firmicutes and Proteobacteria have also been described as the most dominant phyla in the ecosystems of cooling towers and water within sugar-cane processing plants (Sharmin et al., 2013). In fact, most of the identified strains such as *Pseudomonas* sp*., Acinetobacter* sp., *Corynebacterium* sp. have been detected from various wastewater environments (Greay et al., 2019; Han et al., 2020). Besides, *Providencia* sp. was isolated from wastewater and applied to perform microbial remediation (Abo-Amer et al., 2013). A commonly found species, *Legionella,* was not detected in these biofilm samples. *Legionella* has been frequently observed in biofilms obtained from cooling water suspensions (Edagawa et al., 2008; Pereira et al., 2017). Nevertheless, a previous study uncovered that *Pseudomonas* has a robust negative correlation with *Legionella* in cooling towers (Paranjape et al., 2020), which may explain the lack of *Legionella* in the present sampled community. Overall, these isolates are likely to have social interaction related to biofilm formation, as they are co-existence in such an environment.

Building a multi-species biofilm model has more ecological relevance in comparison to individual cultivation model and could serve as an effective way to explore biofilm development in natural conditions (Røder et al., 2016). This is due to the high prevalence of synergistic effects in biofilm formation as observed from different isolates; for example, one bacterial consortium possessed synergy to produce biofilms with the interspecific cooperation (Liu et al., 2019; Ren et al., 2015). In the present work, four species were found to grow individually in the synthetic medium, indicating that they are able to assimilate and tolerate the supplied volatile fatty acids. The latter is essential as these volatile fatty acids can inhibit cell growth or display distinctive toxic effects (Pinhal et al., 2019; Wilbanks and Trinh, 2017). Despite their growth capabilities, only strain P17 was found to have the capacity to form biofilm when cultured alone. *Pseudomonas* species are frequent inhabitants of freshwater environments and colonizers of water supply networks via adhesion and biofilm formation (Pereira et al., 2018). It confirms the vital role of *Pseudomonas* species in the biofilm forming process in the cooling tower. Intriguingly, biofilm biomass of the dual-/ triple- and four species was promoted significantly when P17 was present, showing a synergistic effect between these biofilm production strains. Previous studies have demonstrated that *Pseudomonas* had diverse molecular interactions with *Acinetobacter* sp. (Hansen et al., 2007) and *Corynebacterium* sp. (Brathwaite and Dickey, 1970). Nevertheless, synergic mechanisms among the four species are still not clear, further exploration is needed to reveal the synergy for biofilm formation.

Previous studies have shown that the advantages of metabolic inhibitors in many fields, such as antiparasitic drugs (Mukherjee et al., 2016) and cancer therapy (Ramesh et al., 2020) by blocking specific cellular metabolisms. Many small-molecule agents have recently been developed to treat microbial biofilms based on their mechanistic understanding (Qvortrup et al., 2019). In this study, three metabolic inhibitors were selected as they can target distinct metabolic pathways. Sulfathiazole has been reported to interfere with c-di-GMP biosynthesis and reduce the biofilm formation effectively (Antoniani et al., 2010). 3BP and 3-NP can trap the isocitrate lyase in a catalytic conformation in which the active site is utterly inaccessible to its substrate (Moynihan and Murkin, 2014; Sharma et al., 2000). 3BP and 3-NP are metabolic inhibitors to the enzymes such as succinate dehydrogenase and isocitrate lyase, which are essential to the metabolic pathways for volatile fatty acids catabolism. Isocitrate lyase plays a pivotal role in allowing net carbon gain by diverting acetyl-CoA from β-oxidation of fatty acids into the glyoxylate shunt pathway. 3BP has been also described as a potent inhibitor of glycolysis/gluconeogenesis, which has already been applied to block the energy generation in cancer cells (Darabedian et al., 2018). Indeed, gluconeogenesis is an essential metabolic pathway to convert the intermediates from volatile fatty acids catabolism to precursors of biofilm biomass such as monosaccharides and eDNA. Three inhibitors can impact the cell proliferation and fully inhibit the biofilm formation when used individually, indicating they targeted the essential metabolic pathways by using volatile fatty acids as the carbon sources. The inhibitory efficiency was remarkably improved when using combinations of them, even to the synergistic multi-species biofilm formation. Thus, this is a potent way to prevent cellular metabolism via targeting distinct metabolic pathways by using different inhibitors simultaneously. Unexpectedly, the biofilm formation of P17 and multi-species were promoted under relatively low concentration of the metabolic inhibitors, for instance, treating P17 with ST (16 μg/mL) after 24 h (Fig. 4a), treating multi-species with ST:3BP (16 μg/mL) after 48 h (Fig. 5d). In fact, some antibiotics subinhibitory concentrations have been found to promote microbial biofilm formation by a trade-off between drug toxicity and the beneficial results of cell lysis (Yu et al., 2018). It implies that the trade-off of inhibitor effect and beneficial factors is present when using metabolic inhibitors to prevent biofilm formation. Further inquiry into the interplay between bacterial response and inhibitor efficacy in biofilm development is necessary to determine each inhibitor’s optimal conditions.

## 5. Conclusion

Overall, we isolated the biofilm-forming related microorganisms and developed a multi-species biofilm model by mimicking a natural cooling water system. Culturing results showed that four species were able to assimilate the volatile fatty acids and generate biofilm biomass with a synergistic effect. Combined inhibitors displayed an effective inhibitory effect on the biofilm development, which demonstrated that targeting cellular metabolism is a potent approach to inhibit synergistic biofilm formation of multi-species. Exploring the bacterial responses to these metabolic inhibitors and building more complex biofilm systems will help evaluate their inhibiting efficacy in practice.

## Declaration of competing interest

The authors declare that they have no known competing financial interests or personal relationships that could have appeared to influence the work reported in this paper.

## Acknowledgements

The authors would like to acknowledge Juan Andres Martinez for the language editing and proofreading. This study was supported by Talk2Clean project through the funding from Walloon Region (Grant Number: 7988).

## Notes

### Competing Interest Statement

The authors have declared no competing interest.

