## Supplementary material for "Targeting cellular metabolism to inhibit synergistic biofilm formation of multi-species isolated from a cooling water system"

### (Figure S1 – S5)

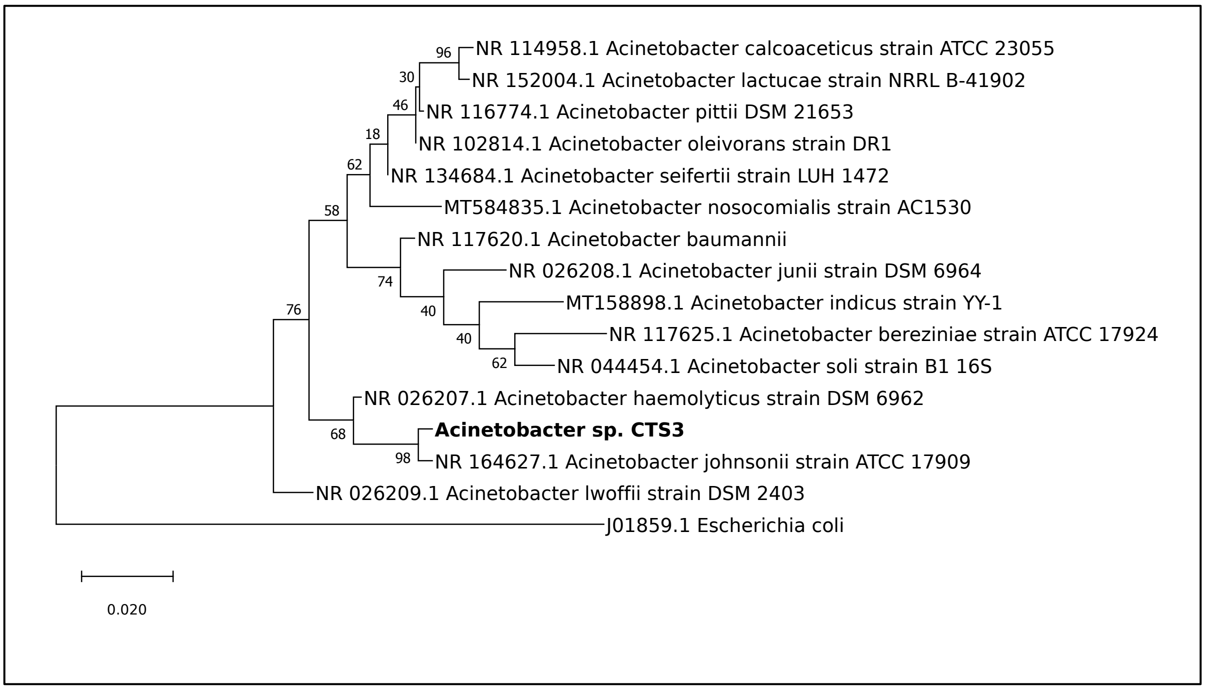


**Fig. S1. Phylogenetic tree of *Acinetobacter* sp. CTS3 (A3).** *Escherichia coli* (Genbank: J01859.1) was set as the outgroup species. Bootstrap values are displayed at each node.

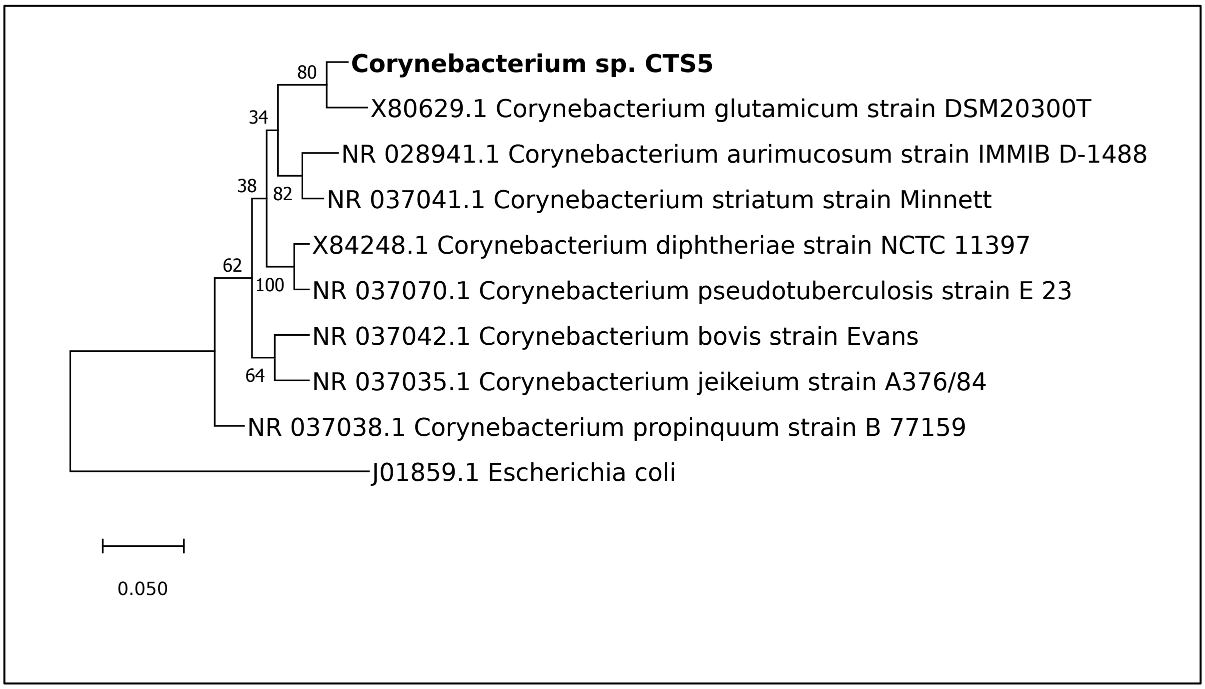


**Fig. S2. Phylogenetic tree of *Corynebacterium* sp. CTS5 (C5).** *Escherichia coli* (Genbank: J01859.1) was set as the outgroup species. Bootstrap values are displayed at each node.

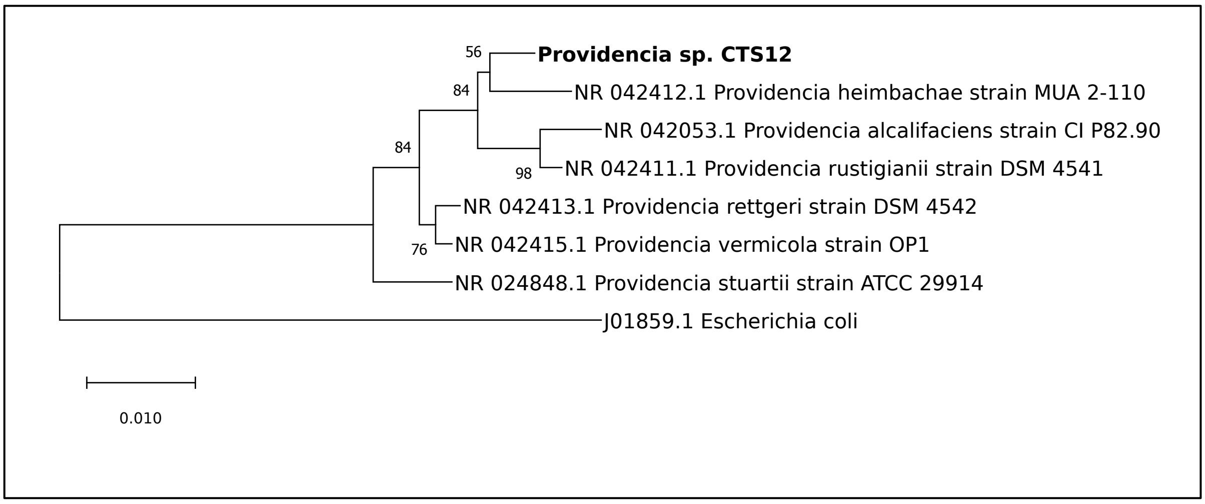


**Fig. S3. Phylogenetic tree of *Providencia* sp. CTS12 (P12).** *Escherichia coli* (Genbank: J01859.1) was set as the outgroup species. Bootstrap values are displayed at each node.

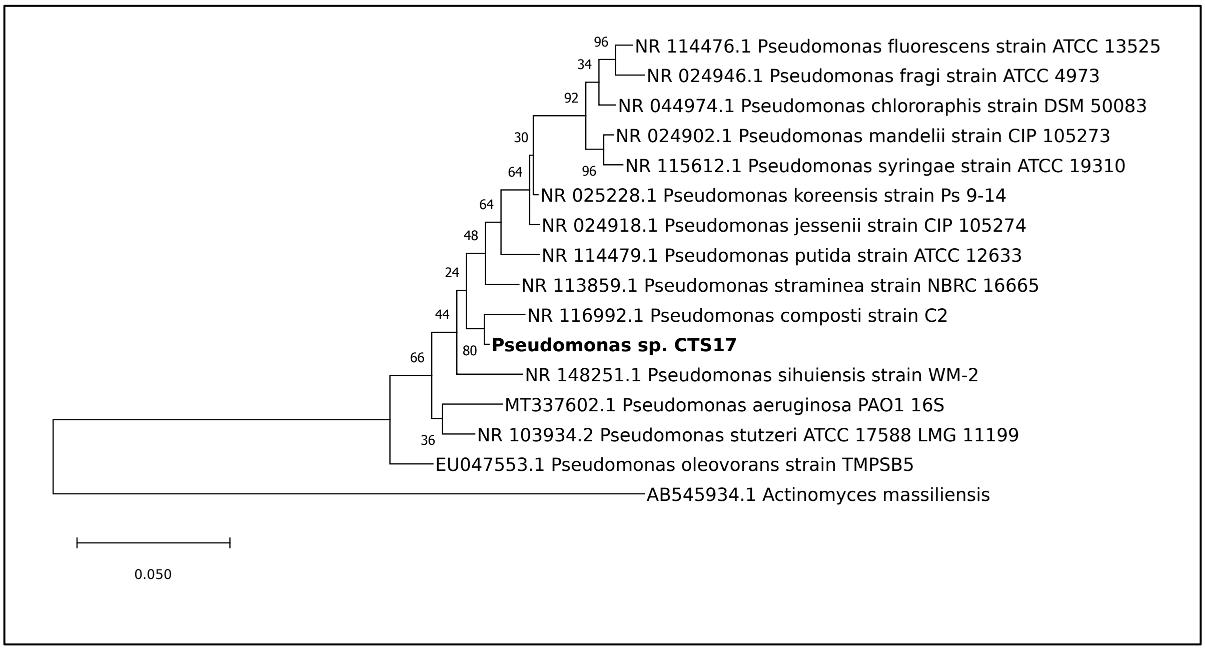


**Fig. S4. Phylogenetic tree of *Pseudomonas* sp. CTS17 (P17).** *Actinomyces massIliensis* was set as the outgroup species. Bootstrap values are displayed at each node.

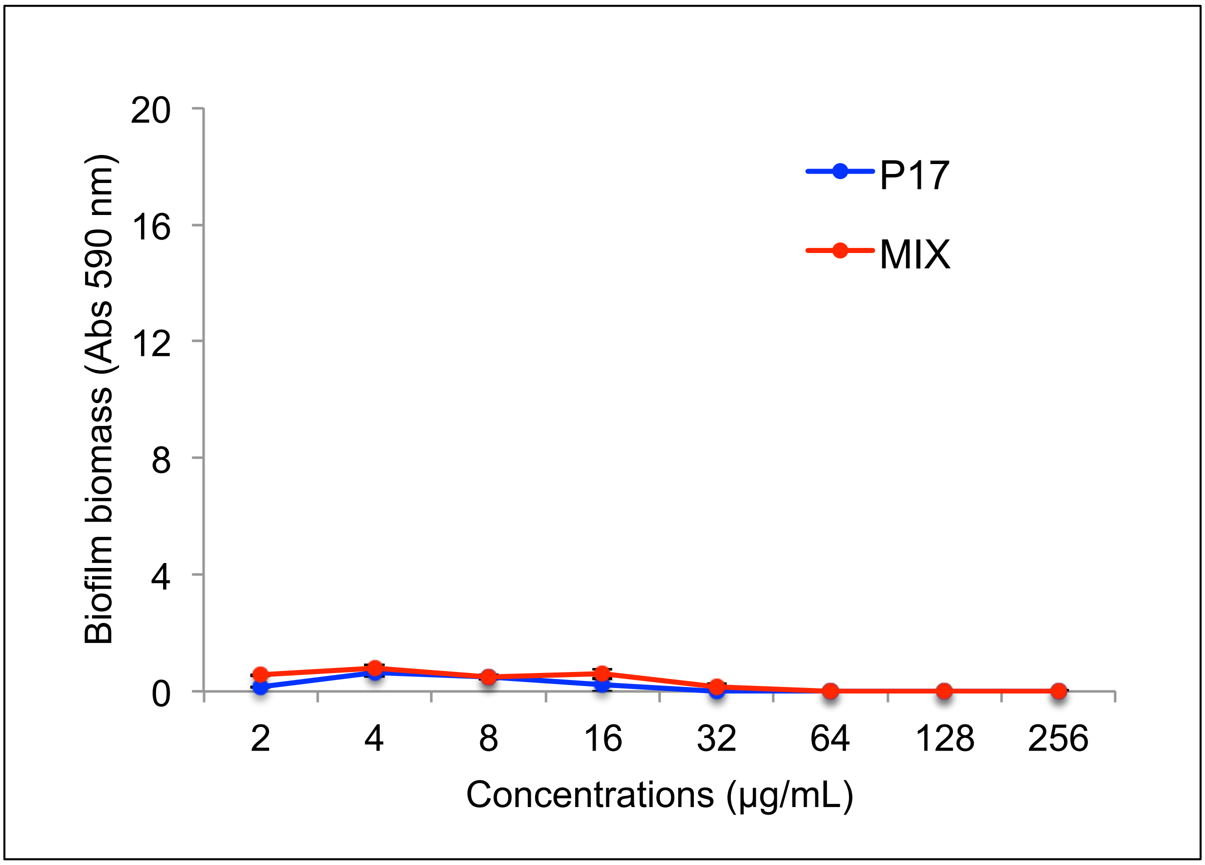


**Fig. S5. Biofilm formation of P17 and four species (MIX) with the treatment of three metabolic inhibitors (ST: 3BP: 3-NP) in the synthetic medium after 96 h, n = 4.**
